# Structural insights into the FtsEX-EnvC complex regulation on septal peptidoglycan hydrolysis in Vibrio cholerae

**DOI:** 10.1101/2023.10.06.561269

**Authors:** Aili Hao, Yang Suo, Seok-Yong Lee

**Affiliations:** Department of Biochemistry, Duke University Medical Center, 303 Research Drive, Durham, North Carolina, 27710, USA

## Abstract

During bacterial cell division, hydrolysis of septal peptidoglycan (sPG) is crucial for cell separation. This sPG hydrolysis is performed by the enzyme amidases whose activity is regulated by the integral membrane protein complex FtsEX-EnvC. FtsEX is an ATP-binding cassette transporter and EnvC is a long coiled-coiled protein that interacts with and activates the amidases. The molecular mechanism by which the FtsEX-EnvC complex activates amidases remains largely unclear. We present the cryo-electron microscopy structure of the FtsEX-EnvC complex from the pathogenic bacteria *V. cholerae* (FtsEX-EnvC_VC_). FtsEX-EnvC_VC_ in the presence of ADP adopts a distinct conformation where EnvC is “horizontally extended” rather than “vertically extended”. Subsequent structural studies suggest that EnvC can swing between these conformations in space in a nucleotide-dependent manner. Our structural analysis and functional studies suggest that FtsEX-EnvCvc may employ spatial control of EnvC for amidase activation, suggesting species-specific diversity in the FtsEX-EnvC regulation on septal peptidoglycan hydrolysis.

## INTRODUCTION

Bacterial cell wall defines the cell shape and protects the cells against extreme osmotic pressures. Bacterial cell wall is made of peptidoglycan (PG). PG is composed of glycan strands with alternating units of N-acetylglucosamine (GlcNAc) and N-acetylmuramic acid (MurNAc), with a pentapeptide stem attached to the MurNAc unit. The glycan polymers are crosslinked via peptide stems, resulting in a continuous mesh-like structure that surrounds the cell ^1^. PG is dynamic; expanding and being cleaved throughout the cell’s life cycle. This PG remodeling play crucial roles in cell growth, separation, and spore maturation^2,3^.

During bacterial cell division, PG layers are formed between the invaginating bilayers of the cytoplasmic membrane, known as the septal PG (sPG). The cell synthesizes new sPG during cell division but also has to hydrolyze it to allow outer membrane constriction^4,5^. This hydrolysis, carried out by LytC type N-acetylmuramyl-L-alanine amidases (amidases), is crucial for outer membrane invagination and daughter cell separation^6–8^.

*E.coli* encodes three amidases: AmiA, AmiB, and AmiC^7,9,10^. These are largely functionally redundant; mutations in all three amidases are required to disrupt daughter cell separation^11^. On the contrary, the enteric pathogen *Vibrio cholerae* encodes only a single amidase^12^. The activity of amidases is tightly regulated, as uncontrolled PG hydrolysis can lead to cell lysis. Amidases require activation by activators containing the LytM domains (also known as peptidase_M23)^13–15^. In both *E.coli* and *V.cholerae* there are more than one LytM-containing factor including EnvC and NlpD^12,14,16^. EnvC from *E. coli* has been demonstrated to stimulate PG hydrolysis activity of AmiA and AmiB *in vitro* ^14^. Although the cellular components for PG remodeling appear to be generally conserved between *E. coli* and *V. cholerae*, their regulatory networks are diverse ^12^.

Notably, EnvC interacts with FtsEX, forming the FtsEX-EnvC complex^17–19^. FtsEX is a component of the divisome, the multiprotein complex responsible for PG synthesis and remodeling during cell division. FtsEX belongs to ATP-binding cassette (ABC) transporter superfamily. Within the ABC transporter superfamily, FtsEX is a member of the type VII subfamily, which includes MacB (PDB: 5LIL) and LolCDE (PBD: 7ARI)^20–22^. This subfamily does not transport substrates. Instead, they utilize ATP hydrolysis to induce conformational changes to carry out mechanical work ^23^. In *E. coli*, the ATPase activity of FtsEX is thought to be required for proper cell division, as an ATPase mutant inhibits cell division^8,23,24^. However, the molecular mechanism of ATP hydrolysis in the FtsEX-EnvC mediated amidase activation remain unclear. Besides regulating amidase activation, FtsEX also promotes cell constriction by interacting with other components of the divisome, which are crucial for cell division ^24–27^.

A recent cryo-EM study of FtsEX-EnvC from *Pseudomonas aeruginosa* unveiled the architecture of the full length FtsEX-EnvC complex^28^. Together with a crystallographic study of FtsX fragment in complex with EnvC from *E. coli* ^29^, these studies provided a model of the conformational change of FtsEX-EnvC leading to amidase activation ^28^.

In our study, we present a structure of the FtsEX-EnvC complex with ADP bound from *V. cholerae* revealing a novel conformation where EnvC is “horizontally extended”. We also obtained a low-resolution single-particle cryo-EM reconstruction of the mutant complex in the absence of nucleotide, displaying a distinct EnvC conformation. This suggests a potential nucleotide dependent motion of EnvC. Based on our structural analysis and functional studies, we proposed a model of amidase activation via FtsEX-EnvC from *V. cholerae*. Our model suggests that different bacterial species might employ diverse mechanism to utilize FtsEX.

## RESULTS

### Structure determination of FtsEX-EnvC and FtsEX

*V. cholerae* FtsE, FtsX and EnvC (FtsEX-EnvC_VC_) were co-expressed in *E.coli* strain C41(DE3). They were purified using lauryl maltose neopentyl glycol (LMNG) and cholesteryl hemi-succinate (CHS). Based on size exclusion chromatography, the three proteins formed a complex (Figure S1A). Details of the purification and cryo-EM sample preparation are provided in the STAR methods.

We incubated the purified complex with ADP•Mg prior to grid preparation. Data were collected using a Titan Krios microscope. Data processing revealed two distinct 3D classes: FtsEX and FtsEX-EnvC. We subsequently resolved 3D reconstructions for FtsEX and FtsEX-EnvC with resolutions of 3.51Å and 3.47Å, respectively (Figure 1A). Data collection, refinement, and validation statistic are provided in Table S1, and the data processing scheme is shown in Figure S2. The cryo-EM density for ADP•Mg was evident in both reconstructions (Figure S3 and S4).

**Figure 1.**
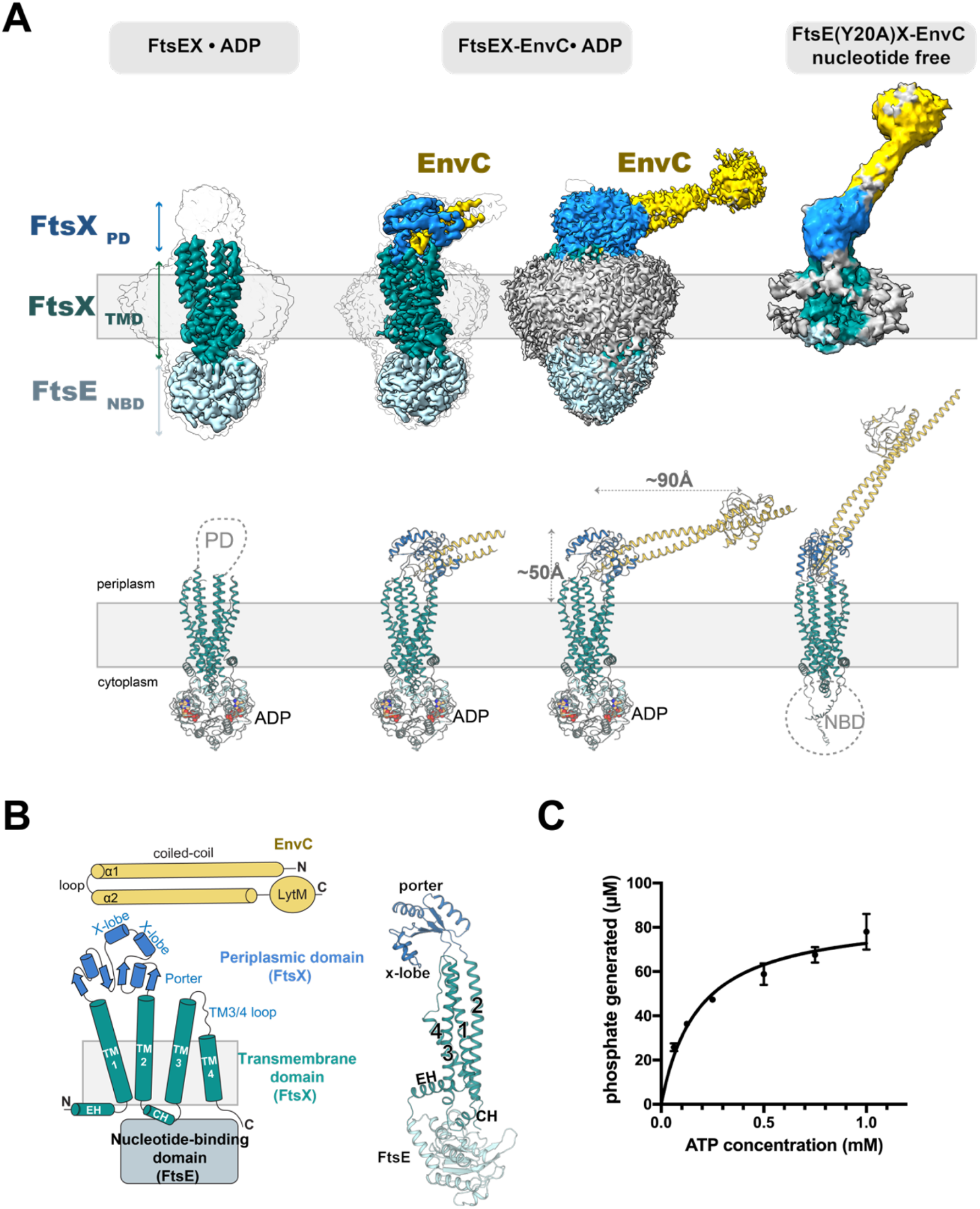
Biochemical characterization and cryo-EM studies of the FtsEX-EnvC_VC_ complex. (**A**) Cryo-EM density maps (top) of FtsEX (ADP bound), FtsEX-EnvC (ADP bound) in two different contour levels, and FtsE(Y20A)X-EnvC (nucleotide-free). The extended region of EnvC is visible at lower contours. The bottom shows the corresponding models in cartoon representation. An AlphaFold predicted model was used for FtsE(Y20A)X-EnvC (nucleotide-free). The unresolvable regions in the cryo-EM reconstructions are shown as dotted lines. (**B**) Topology diagram of the FtsEX-EnvC complex. FtsE is in light cyan; FtsX is divided into the transmembrane domain (TMD, in teal) and the periplasmic domain (PD, in blue); EnvC is colored gold. The color scheme is consistent throughout the figures. (**C**) ATPase activity of FtsEX-EnvC under varying substrate ATP concentrations. Each point represents mean ± s.d.

FtsEX_VC_ is composed of two FtsE and two FtsX. Two FtsE subunits forms the homodimeric nucleotide-binding domain (NBD), while the two FtsX subunits comprise the transmembrane domain (TMD) and the periplasmic domain (PD) (Figure 1B). Each FtsX dimer contains four TM segments (TM1-TM4). TM1 has an L-shape and includes an elbow helix (EH) that runs parallel to the membrane. TM1, TM2 and TM3 are long and extend outside the membrane, while TM4 is a short helix fully embedded within the membrane (Figure 1B). TM1 and TM2 connect to a large periplasmic domain, which is composed of both porter and a X-lobe regions ^29^ (Figure 1B). The interaction between FtsX and FtsE is primarily mediated through a coupling helix (CH) that runs between TM2 and TM3. The purified FtsEX-EnvC_VC_ exhibits ATP hydrolysis and has a Km value of 0.18 mM (Figure 1C).

In our FtsEX_VC_ reconstruction, the cryo-EM density for the PD is weak, suggesting flexibility in the PD. However, in the FtsEX-EnvC_VC_ reconstruction, the PD become ordered, revealing its architecture and interactions with EnvC (Figure 1A). EnvC is composed of two long helices (α1 and α2) separated by a loop, forming a coiled coil. A globular LytM domain is located at the C-terminal part of α2 (Figure 1B). One end of the EnvC coiled coil comprised of α1, loop, α2 is inserted between the two PD of FtsX (Figures 1A, 1B, 2A). In our cryo-EM map, the density for the other end of the coil-coil including the LytM domain of EnvC is weak and is only visible at low map contour levels (Figure 1A). We generated two models for the FtsEX-EnvC: the first contains a truncated EnvC where the density is strong (FtsEX-EnvC short) while in the second model, we used AlphaFold to extend the EnvC model, including the remainder of the coiled coil and LytM, which we subsequently docked into the cryo-EM density (FtsEX-EnvC long). Both FtsEX and FtsEX-EnvC have well-resolved NBD and TMD (Figure S3 and S4).

### EnvC binding induces asymmetry in TMD

The TMD in FtsEX displays a pseudo-two-fold symmetry (Figure 2A and 2B). However, the binding of EnvC to FtsEX induces an asymmetric TMD movement, disrupting the symmetry of the TMD in FtsEX. Compared to the TMD of FtsEX, FtsEX-EnvC_VC_ demonstrates that the N-terminal part of TM2 from protomer 2 rotates outward away from its symmetry axis, resulting in a 7.9 Å displacement (Figure 2B). Similarly, the C-terminal part of TM1 from protomer 1 rotates outward, leading to a 7.3Å displacement. TM3 and TM4 display more subtle movements. Superposition of the two FtsX protomers reveals that, in addition to helical bending, TM2 undergoes secondary structural re-arrangement, as the top of the TM2 in protomer 2 becomes broken and transitions into a loop (Figure 2C). The most prominent symmetry breaking is in the PD (Figure 2C). The two PDs can be superimposed with a Ca R.M.S.D. of 0.99, but they assume distinct orientations – a rotation difference of about 100° (Figure 2C). Since TM1 and TM2 are connected to the PD, the binding of EnvC to the PD and its subsequent conformational changes likely lead to the symmetry disruption of the TMD of the FtsEX-EnvC.

**Figure 2.**
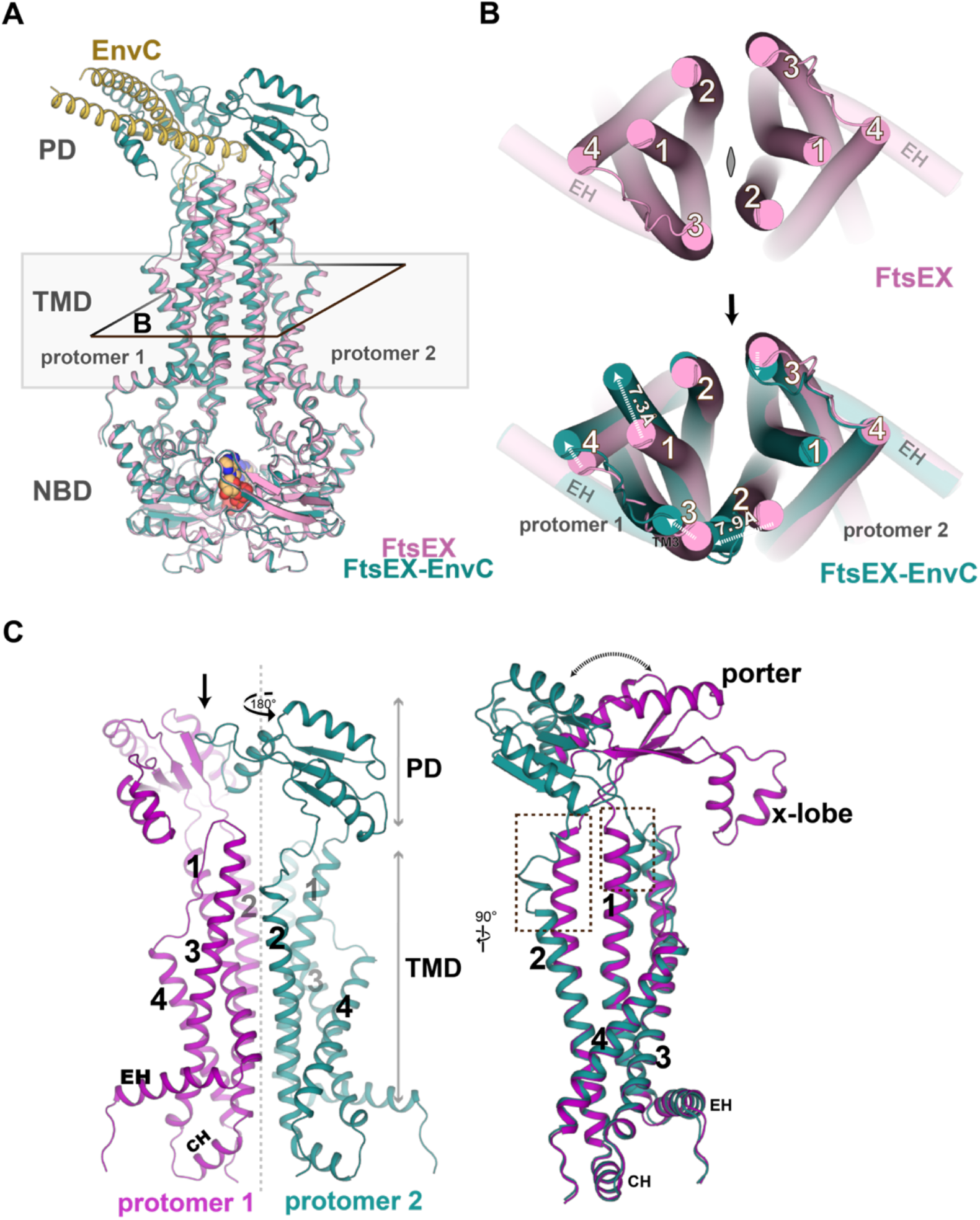
EnvC binding induces asymmetry in FtsEX. (**A**) Overlay of FtsEX and FtsEX-EnvC (both with ADP bound) from the front view (**B**) Top shows the symmetric TMD arrangement in FtsEX. Bottom shows the overlay of TMs arrangements in FtsEX (pink) and FtsEX-EnvC (teal), demonstrating how symmetry is broken upon EnvC binding. White arrow indicates movement in TMs. (**C**) Superposition of the two FtsX protomers from FtsEX-EnvC. Protomer 2 (purple) is rotated 180° to overlay with protomer 1. The periplasmic domains of the two protomers adopt different conformations and are rotated. TM1 and TM2 show movements and conformation differences, highlighted in dotted boxes.

### EnvC adopts a “horizontally extended” conformation in the ADP-bound state

Notably, in our ADP bound FtxEX-EnvC reconstruction, EnvC is oriented nearly parallel to the membrane plane, extending more than 90 Å in length horizontally and ∼50 Å in height from the membrane. We have termed this EnvC conformation as the “horizontally extended” conformation (Figure 1A). This orientation is distinct from that of the recent published cryo-EM structures of FtsEX-EnvC from *P.aeruginosa*^28^ where EnvC is extend vertically into the periplasmic space (Figure S5A).

In our ADP bound FtsEX-EnvC_VC_ structure, extensive interactions between EnvC and FtsX stabilize this horizontally extended conformation. EnvC interacts with FtsX primarily through its two PDs. The PD is divided into the porter and X-lobe regions (Figure 1B). The porter and X-lobe are arranged in a manner resembling a clam shell (Figure 3A). One side of EnvC coiled-coil (α1, loop, α2) is nestled between two X-lobes of the PDs (Figure 3A).

**Figure 3.**
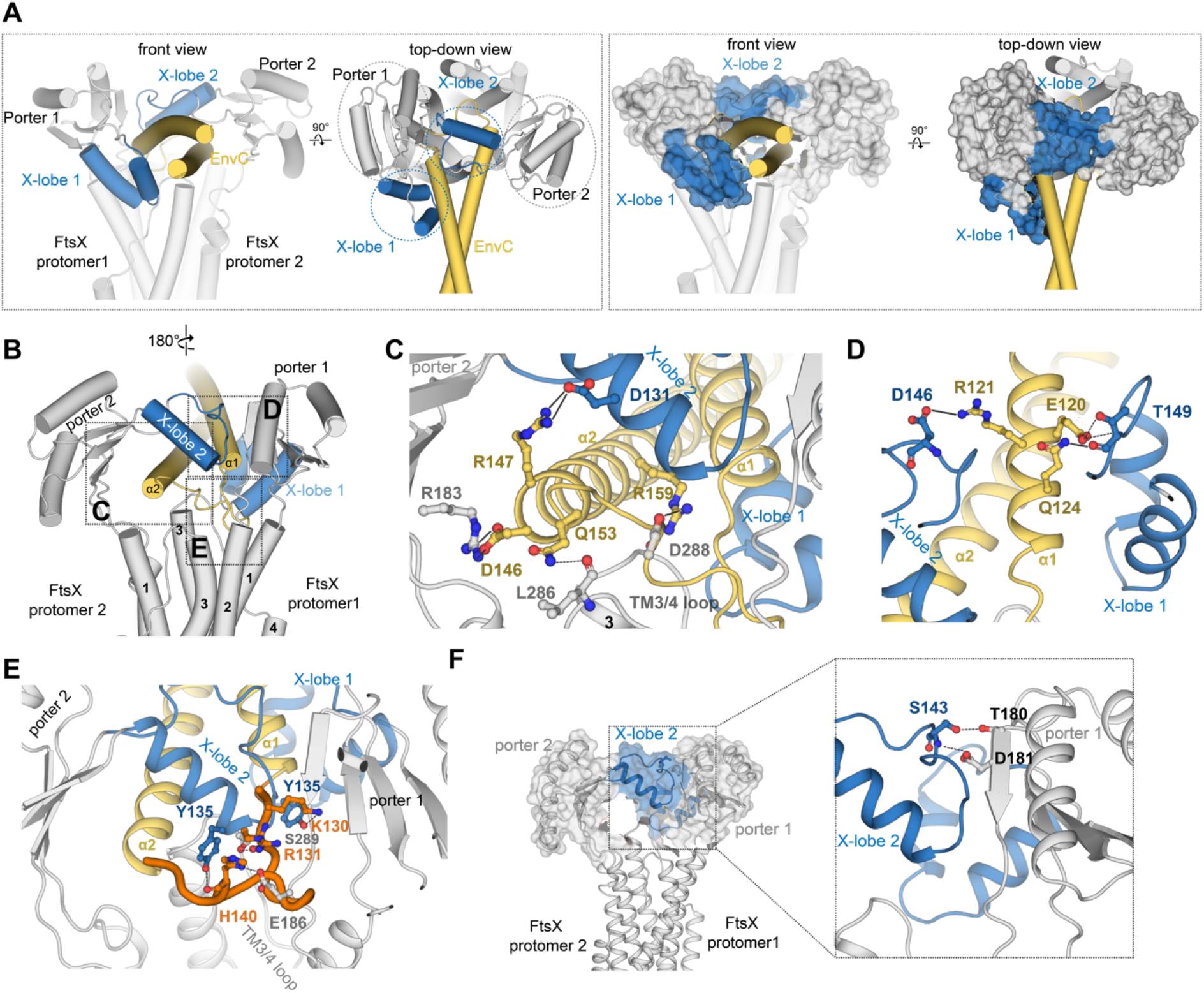
EnvC is sandwiched between two x-lobes. (**A**) Left shows the cartoon representation of the FtsX-EnvC interaction. EnvC is positioned between an X-lobe from each periplasmic domain. Right shows the same views in surface representation. X-lobe is in blue while the porter domain and TMD are in grey. EnvC is in yellow. (**B**) View from the back. Three interaction sites are highlighted in dotted boxes: the α1 helix, the α2 helix, and the loop connecting the two helices. The interaction between FtsX and EnvC is primarily mediated through hydrophobic residues. (**C**) The hydrogen bonds and salt bridges between FtsX and α2 of EnvC. α2 forms hydrogen bonds with a residue from X-lobe 1 (on top), porter 1 (on the left), and the TM3/4 loop (at the bottom). Salt bridges are shown with solid line; hydrogen bonds are shown with dotted lines. (**D**) The hydrogen bonds and salt bridges between FtsX and α1 of EnvC. α1 forms hydrogen bonds with residues from X-lobe 1 (from the left), and X-lobe 2 (from the right) (**E**) The hydrogen bonds and salt bridges between FtsX and the loop of EnvC. The loop interacts with Y135 from each PD, porter 2 from the front, and the TM3/4 loop from the back. (**F**) The hydrogen bonds between the two FtsX protomers: X-lobe from protomer 1 and porter from protomer 2.

The interactions between the FtsX PDs and EnvC are mediated by extensive hydrophobic interactions as well as hydrogen bond interactions. There are three sites on EnvC that interact with X-lobes: α1, α2 and the loop that connects α1 and α2 of EnvC (Figure 3B). Many hydrophobic residues: L119, L122, L123, Y126, Y127, L128, L133, F138, F139, I148, Y151, Y152, L155, A156) on the EnvC as well as (L132, F138, L144, L145, L150) on the FtsX X-lobe constitute these interaction sites. In addition to hydrophobic interactions, H-bonding and salt bridge interactions at each site stabilize the horizontally extended conformation of EnvC (Figure 3C-3E). Concerning the α2-X-lobe interactions, three pairs of salt bridges are formed: FtsX R183 with EnvC D146, FtsX D131 with EnvC R147, FtsX D288 with EnvC R159) (Figure 3C). There is also a hydrogen bond interaction between EnvC D146 and the backbone carbonyl of L286, which is part of the TM3/4 loop (Figure 3C). For the α1 –X lobe interactions, α1 is in close contact with both X-lobes via salt bridges form between EnvC R121 and D146 from X-lobe 2 as well as between EnvC E120 and EnvC Q124 and T149 from X-lobe 1 (Figure 3D). Regarding the loop-X-lobe interactions, within the loop region, EnvC H140 and K130 forms hydrogen bonding network with two Y135 residues from both X-lobes as well as E186 on the TM3/4 loop (Figure 3E). Backbone carbonyl of EnvC R131 forms hydrogen bonding with S289 from the porter 1.

In this “horizontally extended” conformation, the two PDs make close contact between porter 1 and X-lobe 2 (Figure 3F). Additional hydrogen bonding interactions that contribute to this conformation include the sidechain and backbone nitrogen of S143 from X-lobe 2 interacting with T180 and D181 from porter 1 (Figure 3F).

### Conformational landscape of the EnvC in the FtsEX-EnvC_VC_ complex

The LytM domain at the other end of EnvC coiled coil is hypothesized to interact with amidases for activation. Amidases contains a PG-binding domain and are believed to be localized at the PG layer^30^. However, the peptidoglycan layer is located 100-120 Å above the inner membrane plane, so the LytM domain in this “horizontally extended” conformation is unlikely to reach the PG layer. This suggests an inactive state of the complex that cannot reach the amidases. To understand nucleotide-dependent conformation changes in the FtsEX-EnvC complex, we initially attempted to determine the structure of FtsEX-EnvC_VC_ in the presence of ATP or AMP-PNP, but we ended up obtaining the ADP-bound FtsEX-EnvC_VC_ state. To capture different conformations of the complex, we prepared FtsEX-EnvC_VC_ without nucleotides to observe the nucleotide free state conformation. The 2D classes of the dataset revealed conformational heterogeneity (Figure 4A). Some classes resemble the ADP-bound state where EnvC adopts a “horizontally extended” conformation, while others showed EnvC in a more “upward-titled” conformation. Notably, the NBD (FtsE) density appears fuzzy in this class, suggesting that NBD is flexible in the absence of nucleotides. We attempted to use 2D classes that show a “upward tilted conformation of EnvC” to obtain its 3D reconstruction but was not able to achieve a high-resolution reconstruction.

**Figure 4.**
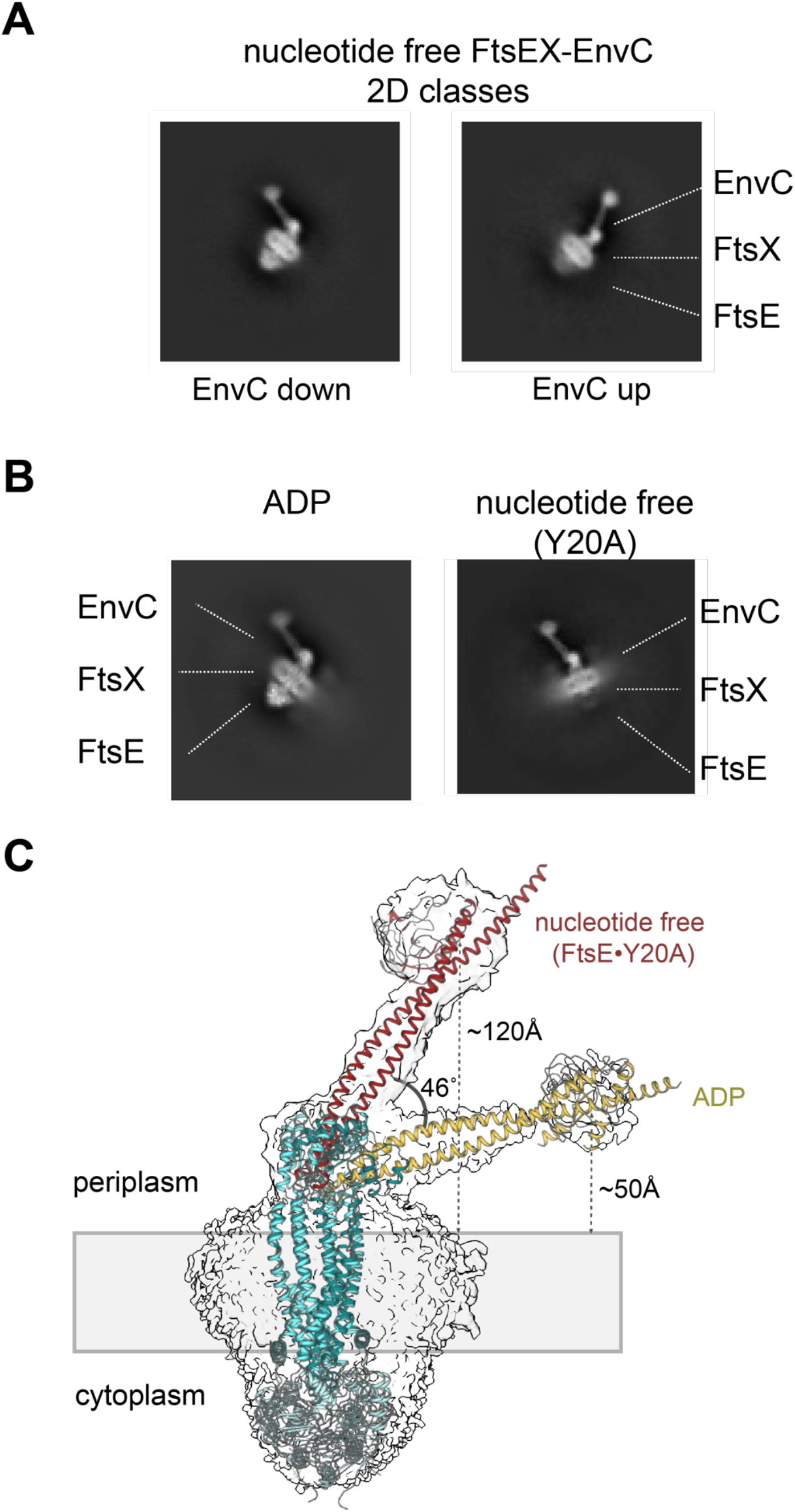
FtsEX-EnvC complex adopts different conformations. (**A**) 2D classification of the nucleotide-free FtsEX-EnvC VC dataset reveals conformational heterogeneity. EnvC can adopt either “horizontally extended” conformation or a “vertically extended” conformation. (**B**) Comparison between the representative 2D class from the ADP dataset and the FtsE(Y20A) mutant dataset. The Y20A mutant shifts the EnvC conformation to almost entirely “vertically extended” conformation. FtsE density in the mutant 2D class appears weak. (**C**) Overlay of the ADP-bound FtsEX-EnvC reconstruction with the nucleotide-free FtsE(Y20A)X-EnvC reconstruction. The model in the FtsE(Y20A)X-EnvC reconstruction is an AlphaFold model docked into the reconstruction. In the nucleotide-free state, EnvC is tilted upward ∼46°, in the nucleotide free state. The LytM domain is positioned ∼110Å away from the membrane in the ADP bound state, whereas it is ∼50Å away in the nucleotide free state.

To increase the population of the complex in the “upward-titled” conformation, we mutated FtsE to favor the complex in a nucleotide free state. We identified sites Y20, K50, D171 and S51 as they are expected to destabilize nucleotide binding and exhibit significantly weaker ATPase activity (Figure S1D). Residue Y20 forms π–π stacking with the nucleotide. Residue K50 is the part of the walker A motif which coordinates with phosphate groups by hydrogen bonding. Residue D171 coordinates with the Mg^2+^ ion. Similarly, S51 also coordinates with Mg^2+^ and the phosphates. We prepared and purified the mutant complexes. Intriguingly, FtsX do not form a stable complex with either FtsE K50A or FtsE D171A mutants indicating that nucleotide binding might be crucial for complex stability (Figure S1B and S1C). We found that FtsE Y20A form a stable complex with FtsX (Figure S1B). Additionally, we found that ATP hydrolysis defective FtsE mutants (Q95A, H204A, E172Q) have no effect on complex formation, suggesting that ATP binding, rather than hydrolysis, might be important for complex stability (Figure S1B).

### Vertically extended conformation of EnvC in FtsE(Y20A)X-EnvC_VC_

We purified the FtsE (Y20A)X-EnvC mutant and determined its structure using single-particle cryo-EM 3D reconstruction. In the 2D classification, all classes showed EnvC in a vertically extended conformation (Figure 4B). Interestingly, we observed that the NBD in the 2D classes appeared blurry, even though they form a stable complex as seen from their size exclusion chromatography profile. This suggests that the NBD might is highly flexible, or it may dissociate from the complex during freezing. For clarity, Figure 4B provides a side-by-side comparison of 2D classes from the wildtype ADP dataset and the Y20A dataset highlighting the different conformations EnvC adopts in different nucleotide states. The 3D reconstruction for the Y20A mutant has a resolution of 7Å. The resulting density of the Y20A mutant on the periplasmic side presents a dumbbell shape; one end is LytM, and the other end the periplasmic domain-EnvC interface. Notably, EnvC adopts a more upright conformation in comparison to its ADP-bound state (Figure 4C). A comparison between the two EnvC conformations suggests a swing motion of ∼46° for EnvC upward, resulting the LytM domain ∼120Å above the membrane plane.

Due to the limited resolution of the 3D reconstruction of FtsE(Y20A)X-EnvC_VC_, we utilized AlphaFold (AF) to generate and dock a model. One model fits the Y20A mutant cryo-EM map well, particularly in the angel between EnvC and FtsX (Figure 4C). We employed this AF model to represent the FtsEX-EnvC_VC_ in the nucleotide-free state and compare it with the structure of ADP bound FtsEX-EnvC_VC_ to gain insight into conformational changes of FtsEX-EnvC_VC_. Since FtsX interacts with EnvC mostly through the two PDs, the drastic rotation movement of EnvC stems from movements in the PDs. Comparing our ADP-bound state model and AF-predicted nucleotide-free model, there is a marked difference in the position of the PD (Figure 5A). Overlaying the PDs from both conformations indicates that one of the PDs (PD2) undergo downward rotation, while the other (PD1) undergoes outward translation as well as downward rotation from the nucleotide-free state to the ADP state (Figure 5B). These movements in the PDs cause EnvC to be repositioned. Since TM1 and TM2 connects the PD, movement in the PD is likely coupled with changes in the TMs (Figure 5C), stemming from the changes in the NBD through the coupling helix. Due to the uncertainty of AF’s predictions for the nucleotide-free conformation of NBD, we avoid a detailed comparison of the NBDs.

**Figure 5.**
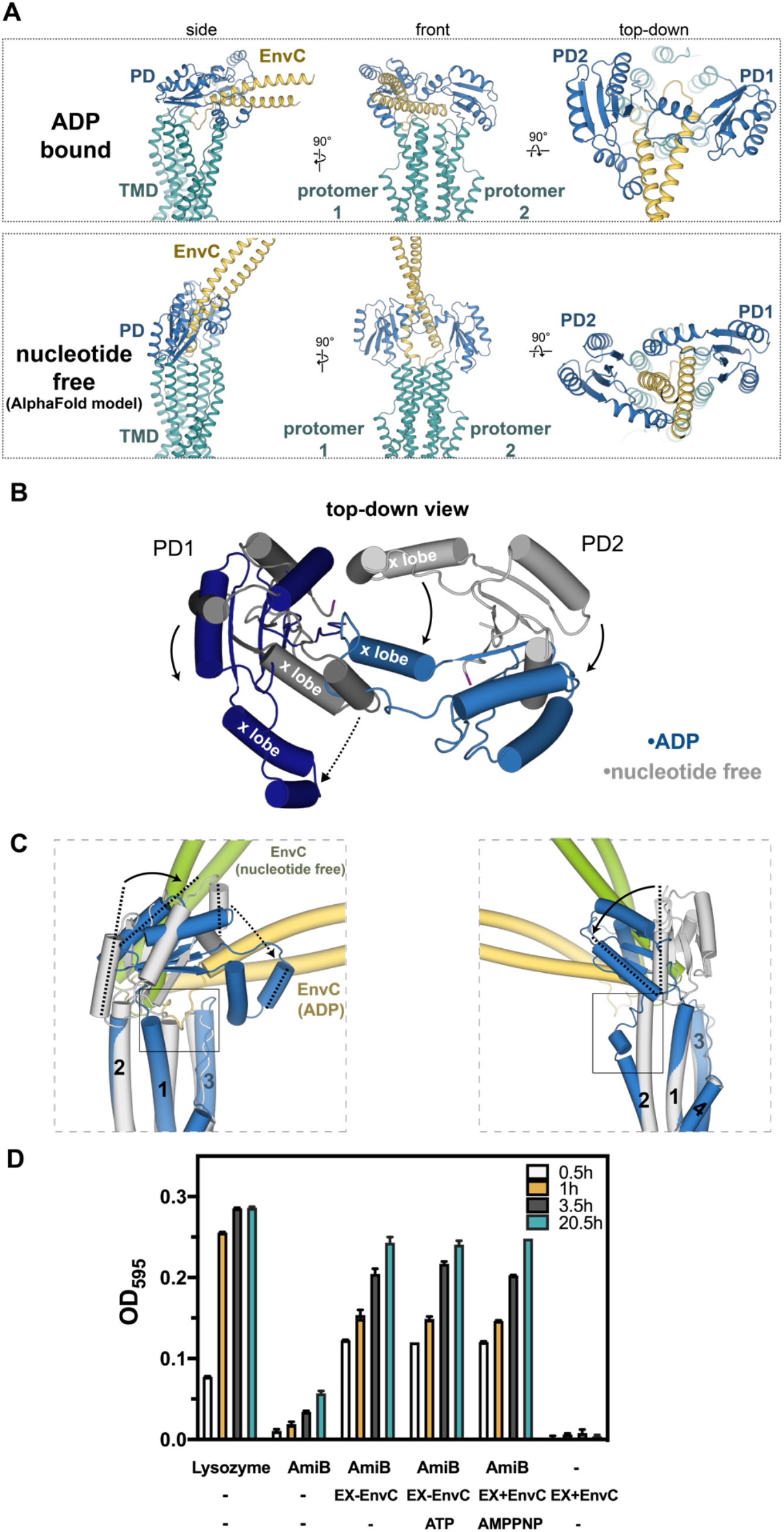
Periplasmic domain movement associated with the EnvC movement. (**A**) Side-by-side comparison of FtsEX-EnvC in the ADP-bound and nucleotide-free states. EnvC is “horizontally extended” in the ADP bound state while it is “vertically extended” in the nucleotide-free state. The PDs from two nucleotide states adopt different conformations. TMD is in teal, PD is in blue and EnvC in gold. (**B**) Detailed comparison of the movement in the PD (only PD are shown). The PD of nucleotide free state is shown in grey, and the PD of ADP bound state is shown in blue. In ADP bound state, PD 1 is rotated downward while PD2 exhibits both a downward rotation and an outward translation. These motions cause EnvC to move downward. (**C**) Zoomed-in view showing the difference in the PD and TM between the two states for each subunit. (**D**) *In vitro* amidase activation of AmiB_VC_ by FtsEX-EnvC_VC_ using the RBB-Dye release assay. Lysozme was used as a positive control. FtsEX-EnvC_VC_ can activate amidase *in vitro* independent of nucleotide presence.

We suggest that FtsEX controls the position of EnvC, providing spatial control over the activation of amidases. To test our spatial control hypothesis, we have performed an *in vitro* amidase activity assay. Our result supports the idea of the spatial regulation on EnvC as no nucleotide is needed for amidase activation in solution due to the absence of spatial separation in this *in vitro* condition (Figure 5D). Both our findings and those reported in the literature clearly show that FtsEX-EnvC or EnvC can activate amidase in solution, without the need for nucleotides^14^.

## DISCUSSION

We report near-atomic structures of FtsEX and FtsEX-EnvC with ADP bound from *V. cholerae*. In the absence of EnvC, the PD of FtsEX is high flexible but is stabilized upon EnvC binding, causing conformational changes at the PD and disrupting the TMD symmetry of FtsEX. The two PDs surround the EnvC through extensive hydrophobic, hydrogen bonding, and salt bridge interactions. Contrary to recent findings on the FtsEX-EnvC complex from *P.auruginosa*, our structure reveals EnvC in a horizontally extended conformation.

Although we have not yet captured the ATP bound state, other recent studies offer insights into its conformation and function. A study detailed the crystal structure of amidase bound to the LytM domain of EnvC, elucidating how LytM activates amidase ^31^. Amidase contains an inhibitory helix blocking its active site, with a parallel regulatory helix. LytM domain interacts with this regulatory helix, displacing the inhibitory helix and exposing the active site. The study posits that ATP binding to FtsX-EnvC induces change that unveil the amidase-binding site.

Another study reported several cryo-EM structures of FtsEX-EnvC from *P. aeruginosa* ^28^. Their structures show EnvC extending into the periplasmic space. Notably, in their ATP-bound FtsE(E163Q)X-EnvC-amidase structure, despite its low resolution, the entire complex spans ∼300Å above the membrane into the periplasmic space. Functional assay suggested that ATP promotes the interaction between AmiB and the FtsEX-EnvC complex ^28^.

Combining our cryo-EM and functional studies with the recent studies, Figure 6 presents our working model for FtsEX-EnvC_VC_. EnvC functions like a long arm that swings up and down under the control of FtsEX. As amidase activation is nucleotide-independent in solution, its suggests that FtsEX from *V. cholerae* primarily regulates the spatial separation between EnvC and the amidases. In the nucleotide free state of FtsEX, EnvC assumes a vertically extended conformation. We speculate that ATP binding induces conformational changes of EnvC to displace the auto-inhibitory helix in amidases. Following ATP hydrolysis, FtsEX repositions the EnvC away from the amidases through a downward rotation of EnvC, reverting the amidases to their auto-inhibited state.

**Figure 6.**
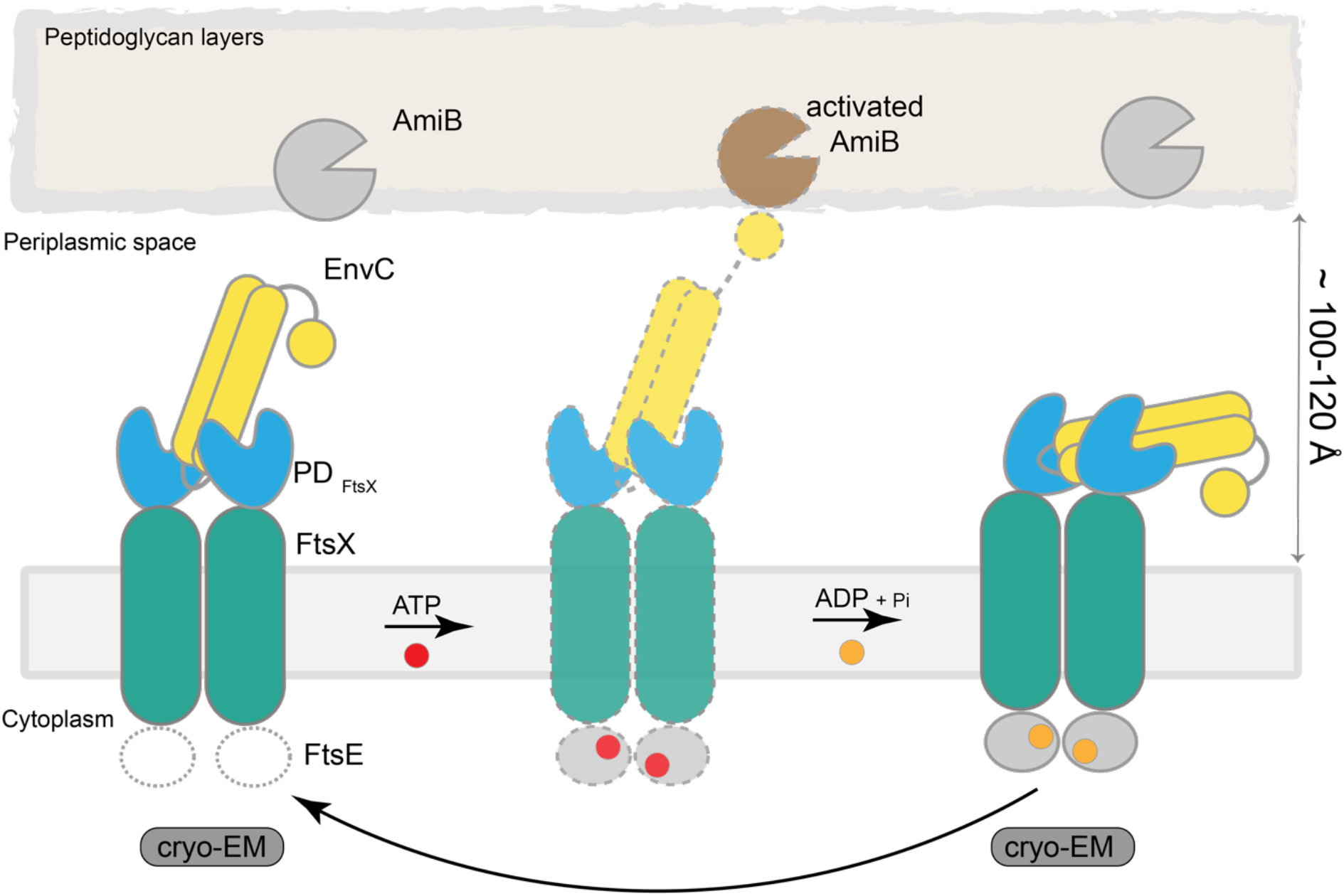
A working model of FtsEX-EnvC in V.cholera. In the nucleotide free state, FtsEX-EnvC adopts a “vertically extended conformation”, as observed in our cryo-EM study. ATP binding likely induces conformational changes in EnvC extending the LytM domain and interacts with amidase, enabling its activation. ATP hydrolysis repositions EnvC into the “horizontally extended” conformation through conformational changes of the PDs. In this state, the LytM domain is distant from the PG layer, returning the amidases to their inhibited state. Dotted line represent hypothetical conformations not captured in cryo-EM.

The observed conformational differences between the cryo-EM structures of *P. aeruginosa* FtsEX-EnvC and our *V.cholera* FtsEX-EnvC structures may arise from species-specific variations, especially considering the diversity in the PG remodeling regulatory network and components are diverse between *E. coli* and *V.cholerae* ^32^. Importantly, the sequences of the X-lobe and the α1-loop-α2 region of EnvC underscore a marked variation between *V.cholera* and *P.aeruginosa* (Figure S5B). Interestingly, while *V. cholera* is characterized as a comma-shaped curved bacterium, *P. aeruginosa* is a regular rod-shaped bacterium (Figure S5A). The difference in cell shape could affect the FtsEX-EnvC mechanism, given its role in dictating the division and the form of the bacteria. An alternate possibility is that our ADP bound conformation of FtsEX-EnvC_VC_ captures a conformational state not previously observed in the conformational landscape of FtsEX-EnvC.

## ACKNOWLEDGMENTS

Cryo-EM data were screened and collected at the Duke University Shared Materials Instrumentation Facility (SMIF). We thank Nilakshee Bhattacharya at SMIF for assistance with the microscope operation. We thank Joe Lutkenhaus and Sebastien Pichoff for performing complementation experiments in E. coli. We thank Ying Yin and Dohoon Kwon for advice throughout the project. This research was supported by Duke Science Technology Scholar Funds (S.-Y.L.). DUKE SMIF is affiliated with the North Carolina Research Triangle Nanotechnology Network, which is in part supported by the NSF (ECCS-2025064).

## AUTHOR CONTRIBUTIONS

A.H. performed all the experiments. Y.S. collected data and helped data processing. A.H. and S.-Y.L. performed model building. S.-Y.L. supervised the experiments. All authors contributed to the analysis of the data and the interpretation of the results. A.H. and S.-Y.L. wrote the manuscript with contributions from Y.S.

## STAR METHODS

## LEAD CONTACT AND MATERIALS AVAILABILITY

Further information and requests for resources and reagents should be directed to and will be fulfilled by the Lead Contact, Seok-Yong Lee.

## EXPERIMENTAL MODEL AND SUBJECT DETAILS

*V. cholera* FtsE, FtsX and EnvC gene are commercially synthesized (BioBasic) and cloned into a modified pCDFDuet vector that contains three T7 promoters. FtsE mutants was generated by QuickChange site-directed mutagenesis kit (Agilent). Recombinant FtsEX-EnvC and mutants proteins were overexpressed in E.coli C41 (DE3) cells in Terrific broth (TB) liquid media. Cells were grown at 37°C at 220rpm until OD reaches 1.0-1.2. Addition of IPTG to a final concentration of 0.4µM initiates induction. Cells were grown overnight ∼18h at 20°C before harvest.

*V.cholera* AmiB gene were synthesized (BioBasic) and cloned into pETDuet vector. The construct contains PPX site and a 10xHis tag at the c-terminus of the gene. Signal peptide (2-34) were removed by QuickChange site-directed mutagenesis kit (Agilent). Recombinant AmiB were overexpressed in *E.coli* C41 (DE3) cells in Terrific broth (TB) liquid media. Cells were grown at 37°C at 220rpm until OD reaches 0.6-0.7. Addition of IPTG to a final concentration of 0.4µM initiates induction. Cells were grown for 4-5 h at 30°C before harvest.

## METHOD DETAILS

### Expression and purification of FtsEX-EnvC

#### Protein expression and purification

To culture and overexpress FtsEX-EnvC complex, the vector contains three T7 promoters and allows expression of three genes. FtsE contains a N-terminal FLAG tag, followed by a PreScission protease (PPX) site cleavage site while FtsX and EnvC have no tags (FtsE-FtsX-EnvC_pCDFDuet). All purification steps were performed at 4 °C. The bacterial cell pellet was collected and resuspended in 50 mM Tris, 150 mM NaCl, 1 μg/mL leupeptin, 1 μg/mL pepstatin, 1 μg/mL aprotinin, DNase I, 1 mM phenylmethylsulphonyl fluoride (PMSF), 10% glycerol. The cells were lysed by sonication. The complex was extracted with 1% LMNG, 0.1% CHS (Anatrace) for 1 hour, followed by centrifugation at 13000 × *g* for 40 min to remove insoluble material. The supernatant was subsequently incubated with FLAG M-2resin (Sigma) for 30min. The resin was collected and washed with 10 column volumes (CV) of wash buffer 50 mM Tris, 500 mM NaCl, 0.1% LMNG, 0.01% CHS, 10% glycerol followed by 10 column volumes of wash buffer 50 mM Tris, 150 mM NaCl, 0.1% LMNG, 0.01% CHS, 10% glycerol. The complex was eluted in the presence of 0.2 mg/ml of FLAG peptide. FtsEX-EnvC was purified by size-exclusion chromatography (SEC) with a Superose 6 10/300 GL column in 20 mM HEPES, 150 mM NaCl, 0.003% LMNG, 0.0003% CHS. The peak fraction(s) containing the FtsEX-EnvC were collected and concentrated to ∼1 mg/mL for cryo-EM sample preparation.

For FtsE(Y20A)X-EnvC mutant, similar protocol was carried out for purification. The sample was collected and concentrated to ∼2mg/ml for cryo-EM sample preparation.

For AmiB(ΔSignal Peptide) expression and purification, the bacterial cell pellet was collected and resuspended in 50 mM Tris, 150 mM NaCl, 10% glycerol and 1% PMSF. The cells were lysed by sonication, followed by centrifugation at 13000 × *g* for 40 min to remove insoluble material. The supernatant was subsequently incubated with cobalt affinity resin (Talon) for 30min. The resin was collected and washed with 10 column volumes (CV) of wash buffer 50 mM Tris, 500 mM NaCl, 10% glycerol, 15mM Imidazole, followed by 10 column volumes of wash buffer 50 mM Tris, 150 mM NaCl, 10% glycerol, 15mM imidazole. The complex was eluted in the presence of 200mM imidazole. FtsEX-EnvC was purified by size-exclusion chromatography with a Superdex 200 10/300 GL column in 20 mM HEPES, 150 mM NaCl. The peak fraction(s) containing AmiB were collected and concentrated. 20% final concentration of glycerol were added to the sample and flash-frozen and store at –80°C until use.

#### Cryo-EM sample preparation

The protein (wildtype or Y20A mutant) was pegylated with MS(PEG)12 methyl-PEG-NHS-ester (Thermo Fisher) at 1:30 or 1:60 molar ratio on ice for 2 hours and then proceed to grid freezing. For FtsEX-EnvC, sample was then incubated with 5mM ADP and 2mM of MgCl2 for at least 30 min on ice before freezing. 2% DMSO was added to the samples right before freezing. The samples were plunge frozen using Leica EM GP2 Automatic Plunger Freezer at 4 °C and 90% humidity. A sample volume of 3 μL sample was applied to a freshly glow-discharged UltrAuFoil R1.2/1.3 300 mesh (Quantifoil); the sample was incubated for 1min and blotted with Whatman No. 1 filter paper for 3 s followed by plunge-freezing in liquid ethane cooled by liquid nitrogen. FtsEX-EnvC•ADP and FtE(Y20A)X-EnvC datasets were collected using a Titan Krios (Thermo Fisher) transmission electron microscope operating at 300 kV equipped with a K3 detector (Gatan) in counting mode with a BioQuantum GIF energy filter (slit width of 20 eV), using the Latitude-S (Gatan) single-particle data acquisition program. Data were collected at a magnification of ×81,000 with a pixel size of 1.08 Å at the specimen level.

For FtsEX-EnvC•ADP dataset, 5091 movies were collected, and each movie contained 60 frames over 4.6 s exposure time, using a dose rate of 15 e-/pix/s for a total accumulated dose of ∼60 e-/Å^2^. The nominal defocus range was set from –0.8 to –1.8 μm. For FtsE(Y20A)X-EnvC dataset, 4590 movies were collected and each movie contained 60 frames over 4.6 s exposure time, using a dose rate of 15 e-/pix/s for a total accumulated dose of ∼60 e-/Å^2^. The nominal defocus range was set from –0.8 to –1.8 μm.

### Cryo-EM data processing

#### FtsEX and FtsEX-EnvC

Beam-induced motion correction and dose-weighing was performed using MotionCor2^1^, followed by CTF estimation using Gctf ^2^. Particle picking was performed with Gaussian picking in RELION^3^. 2,434,011 particles were picked and then extracted with a 64-pixel 4x binned box size at 4.32 Å per pixel and subjected to two-dimensional (2D) classification in cryoSPARC^4^. A total of 537,027 particles were selected and re-extracted unbin at 1.08 Å per pixel which then subjected to cryoSPARC 1-class ab-initio reconstruction to generate a reference. The particles were transfer back to RELION and subjected to 3D classification. One of the classes showed low resolution features which were discarded. The other two classes showed high resolution features while one showed clear strong density at EnvC. The two classes were then processed separately. For class with strong EnvC density, it was subjected to Bayesian polishing, followed by non-uniform refinement^5^ and local refinement in cryoSPARC. The particles were then re-extract to 450-pixel box size and more density showed up. A mask was generated to over the EnvC and PD domain for signal subtraction to boost resolution at that region. The best 179,239 particles were selected and reverted. The particles were then subjected to another round of Bayesian polishing and transferred to cryoSPARC for non-uniform refinement and local refinement, which yielded a reconstruction to 2.55-Å resolution (Figure S2). Map were sharpened with a B-factor of –50.

For class with weak EnvC density, the particles were subjected to another round of 3D classification and the highest resolution class (261, 627 particles) were selected. These particles were subject to one round of Bayesian polishing, followed by non-uniform refinement and local refinement in cryoSPARC, which yield a 3.51-Å resolution reconstruction (Figure S2). Since this class show no EnvC density, it was assigned as FtsEX class.

FtsE(Y20A)X-EnvC: Beam-induced motion correction and dose-weighing was performed using MotionCor2^1^. Corrected micrographs were then imported into cryoSPARC for contrast transfer function (CTF) estimation with patch CTF^4^. Particle picking was performed with blob picker in cryoSPARC. Particles were then extracted with a 64-pixel 4x binned box size at 4.32 Å per pixel and subjected to two-dimensional (2D) classification. A total of 593,477 particles were selected and re-extracted to 512pix box at 1.08 Å per pixel which then subjected to cryoSPARC 2-class ab-initio. 341,344 particles were selected and subjected to non-uniform refinement, which yielded a reconstruction to ∼7.3 Å resolution (Figure S2).

### Model building, refinement and alignment

FtsEX-EnvC short: The previously published LolCDE structure (PDB 7ARL) was used as a reference for FtsE and TMD region for FtsEX-EnvC model building. The previously published E.coli EnvC-periplasmic FtsX structure (PDB 6TPI) were used as a references for PD and EnvC region. During model building the register assignment was guided by the presence of large aromatic side chains. EnvC was truncated to residue 91-191 due to weak density at extended region.

The structures were then manually refined using real-space refinement in Coot with ideal geometry restraints^6^. The restraints for ADP were calculated in Elbow (as implemented in Phenix^7^) from isomeric SMILES strings. The MolProbity^8^ server (http://molprobity.biochem.duke.edu) was utilized to identify problematic regions in the models, which were then manually adjusted in Coot. The final refinement was performed using the phenix-real_space_refine function with global minimization and secondary structure restraints as implemented in the Phenix suite^7^. Structural analyses and illustrations were performed using PYMOL (Schrödinger) and UCSF Chimera X^9^. Data collection and refinement statistics are provided in Table S1.

FtsEX-EnvC long: We used AlphaFold predicted V. cholera EnvC to docked into the cryo-EM density, extending the FtsEX-EnvC short model to generate the FtsEX-EnvC long model. The MolProbity server (http://molprobity.biochem.duke.edu) was utilized to identify problematic regions in the models, which were then manually adjusted in Coot. The final refinement was performed using the phenix-real_space_refine function with global minimization and secondary structure restraints as implemented in the Phenix suite. Structural analyses and illustrations were performed using PYMOL (Schrödinger) and UCSF Chimera X67 FtsEX: FtsEX-EnvC short model was used as a reference. The structures were then manually refined using real-space refinement in Coot with ideal geometry restraints^6^. The restraints for ADP were calculated in Elbow (as implemented in Phenix^7^) from isomeric SMILES strings. The MolProbity server (http://molprobity.biochem.duke.edu) was utilized to identify problematic regions in the models, which were then manually adjusted in Coot. The final refinement was performed using the phenix-real_space_refine function with global minimization and secondary structure restraints as implemented in the Phenix suite. Structural analyses and illustrations were performed using PYMOL (Schrödinger) and UCSF Chimera X. Data collection and refinement statistics are provided in Table 1S.

### Purification and RBB labelling of PG

The purification of PG protocol is adopted from^10–12^. *E.coli* cells (any strain) were grown to late exponential growth stage which is OD∼0.6-1. The cells were harvested and resuspended in PBS. The cells were boiled in 5% of SDS for 30 min at room temperature with vigorous stirring and continuing the stirring overnight. The solution was then spun at 100,000 x g for 1 hour at room temperature to sediment the sacculi. The isolated sacculi were then washed with water 3 times to remove SDS by spin at 100,000 x g. The isolated clean sacculi were then homogenized in PBS and treated with 300 μg/ml α-Chymotrypsin at 37 C° overnight. Another 300 μg/ml α-Chymotrypsin were added the next morning for at least 3 hours. 5% SDS was added and then washed away by centrifugation to remove the supernatant. The α-Chymotrypsin treated sacculi were then incubated with 20 mM RBB (Sigma) in 0.25 M NaOH overnight at 37 °C. The sample was neutralized with HCl, and RBB-labelled sacculi were pelleted by centrifugation (21000 x g, 20min, room temperature). The sacculi were then repeatedly resuspended in water and pelleted by centrifugation until the supernatant was clear. The final pellet was resuspended in water containing 0.02% azide and stored at 4 C° for use.

### *In vitro* amidase RBB-labelled PG hydrolysis assay

The protocol was adapted and modified from Uehara et al. and Rocaboy et al^10,11^. 5 μL of RBB-labelled sacculi were incubated at 37 °C for various times with purified AmiB and/or with FtsEX-EnvC (stored in 50 mM HEPES, pH7.5, 300 mM NaCl, 5% glycerol, 0.005% LMNG, 0.0005% CHS) in a reaction volume of 50 μL. The final protein concentration is 4 μM for AmiB and 1μM FtsEX-EnvC. Lysozyme was used as a positive control at a 4 μM concentration. If nucleotide was added, the nucleotide concentration was 1 mM. Reactions were incubated at 37 C° for various time.

For each time point, all reactions were centrifuged at 18,000 x g for 10 min at room temperature. Supernatants were removed and their absorbance was measured at 595 nm.

### ATPase assay

ATPase activity of FtsEX-EnvC was measured using a malachite green based colorimetric ATPase kit (MAK307, Sigma Aldrich) according to the manufacturer’s instructions. Briefly, 4 μg of FtsEX-EnvC was incubated in 20 mM HEPES pH 7.5, 150mM NaCl, varying concentration of ATP, 1mM MgCl2, 0.01% LMNG, and 0.001% CHS in a reaction volume of 10µL for 30min at 37°C. The reaction was stopped by the addition the malachite green reagent provided in the kit, incubated at room temperature for 30 min and the absorbance at 620 nm was measured using a SpectraMax M spectrophotometer (Molecular Devices). Phosphate standard curve was constructed using the stock solutions provided in the kit and was used to determine the total concentration of released phosphate. Data were plotted and visualized in GraphPad Prism 8.

## DATA AND SOFTWARE AVAILABILITY

The coordinates are deposited in the Protein Data Bank with the PDB ID 8TZJ (FtsEX), 8TZK (FtsEX-EnvC short) and 8TZL (FtsEX-EnvC long). The cryo-EM density maps have been deposited in EMDB with the ID EMD-41760 (FtsEX), EMD-41761 (FtsEX-EnvC) and EMD-41762 (FtsE(Y20A)X-EnvC).

---

**Supplementary Table 1.** Cryo-EM data collection, refinement, and validation statistics for FtsEX, FtsEX-EnvC.

|  | FtsEX<br>ADP•Mg | FtsEX-EnvC<br>short<br>ADP•Mg | FtsEX-EnvC<br>long<br>ADP•Mg |
| --- | --- | --- | --- |
| **Data collection and processing** |  |  |  |
| Magnification | 81,000 | 81,000 | 81,000 |
| Voltage (kV) | 300 | 300 | 300 |
| Electron exposure (e-/Å<sup>2</sup>) | 60 | 60 | 60 |
| Defocus range (μm) | -2 to -1 | -2 to -1 | -2 to -1 |
| Pixel size (Å) | 1.08 | 1.08 | 1.08 |
| Symmetry imposed | C1 | C1 | C1 |
| Initial particle images (no.) | 2,434,011 | 2,434,011 | 2,434,011 |
| Final particle images (no.) | 261,627 | 179,239 | 179,239 |
| Map resolution (Å) | 3.51 | 3.55 | 3.55 |
| FSC threshold | 0.143 | 0.143 | 0.143 |
| **Refinement** |  |  |  |
| Initial model used (PDB code) | FtsEX-EnvC | 7ARL, 6TPI | FtsEX-EnvC short<br>AlphFold EnvCvc |
| Model resolution (Å) | 3.51 | 3.55 | 3.55 |
| FSC threshold | 0.143 | 0.143 | 0.143 |
| Map sharpening B factor (Å<sup>2</sup>) | -50 | -50 | -50 |
| **Model composition** |  |  |  |
| Nonhydrogen atoms | 6574 | 8609 | 9833 |
| Protein residues | 846 | 1149 | 1398 |
| Ligands | 2 | 2 | 2 |
| Water | 0 | 0 | 0 |
| **B factors (Å<sup>2</sup>)** |  |  |  |
| Protein | 74.14 | 83.24 | 142.79 |
| Ligand | 69.98 | 71.69 | 81.31 |
| **R.m.s. deviations** |  |  |  |
| Bond lengths (Å) | 0.002 | 0.003 | 0.003 |
| Bond angles (°) | 0.420 | 0.423 | 0.417 |
| **Validation** |  |  |  |
| MolProbity score | 0.98 | 1.25 | 1.20 |
| Clashscore | 2.1 | 2.81 | 2.49 |
| Poor rotamers (%) | 0 | 0 | 0 |
| **Ramachandran plot** |  |  |  |
| Favored (%) | 96.62 | 96.63 | 96.87 |
| Allowed (%) | 3.38 | 3.37 | 3.13 |
| Disallowed (%) | 0 | 0 | 0 |

**Supplementary Figure 1.**
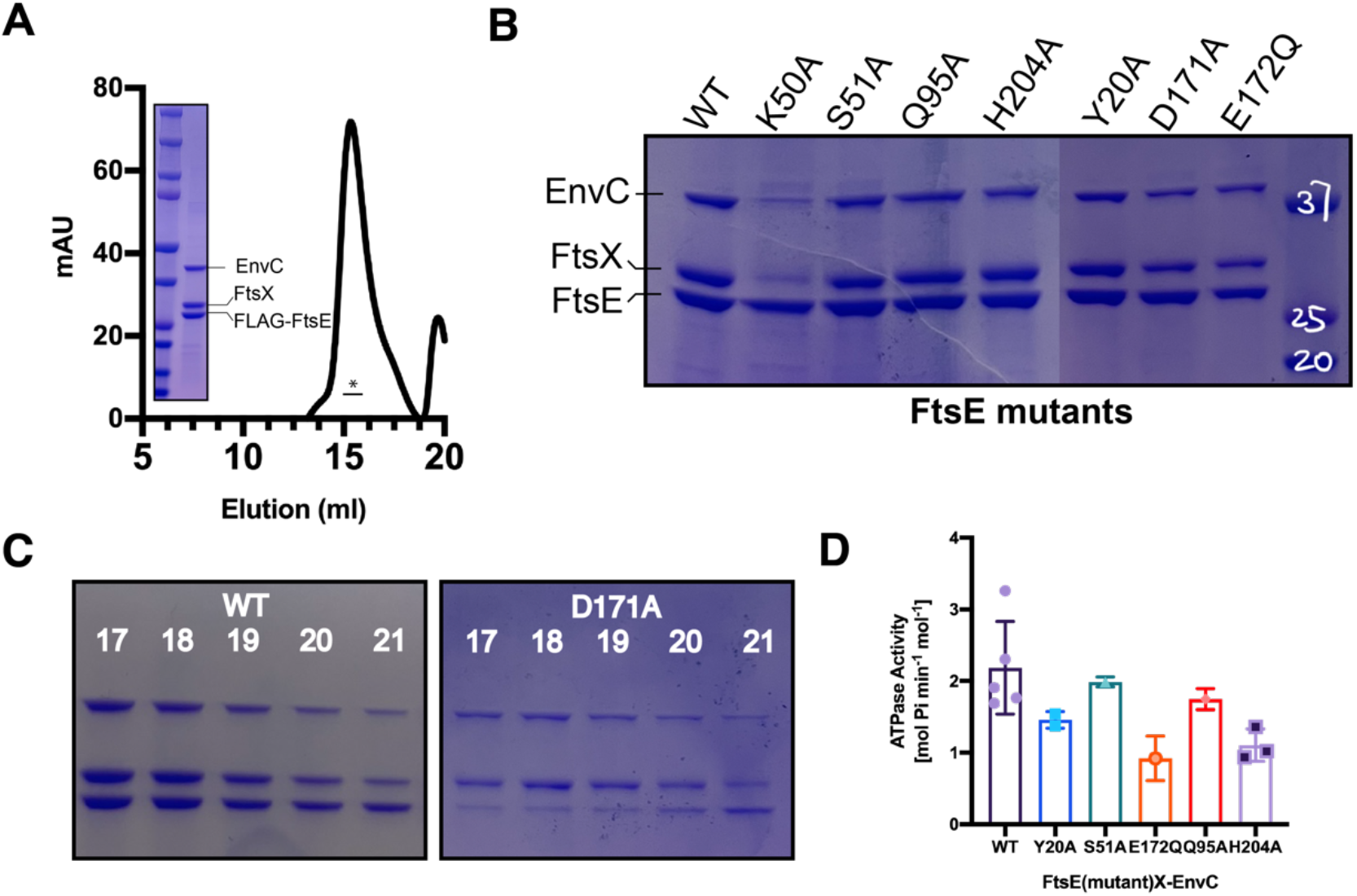
Biochemical characterization of FtsEX-EnvC and mutants. (**A**) SEC profile and SDS-PAGE gel of purified FtsEX-EnvC complex in detergent LMNG/CHS. **(B)** Pull-down of the FtsEX-EnvC complex with different FtsE mutants. The tag is located at the FtsE while FtsX and EnvC has no tag. K50A, a mutation that dis-stablize ATP binding, impairs complex formation. **(C)** Gel filtration fractions of FtsE(WT)X-EnvC and FtsE(D171A)X-EnvC. The D171A mutant complex showed significant dissociation and hence the bands did not co-migrate. D171 is expected to coordinate with Mg so mutation D171A has impaired ability to bind to ATP. **(D)** ATPase activity of FtsEX-EnvC WT or with mutants FtsE.

**Supplementary Figure 2.**
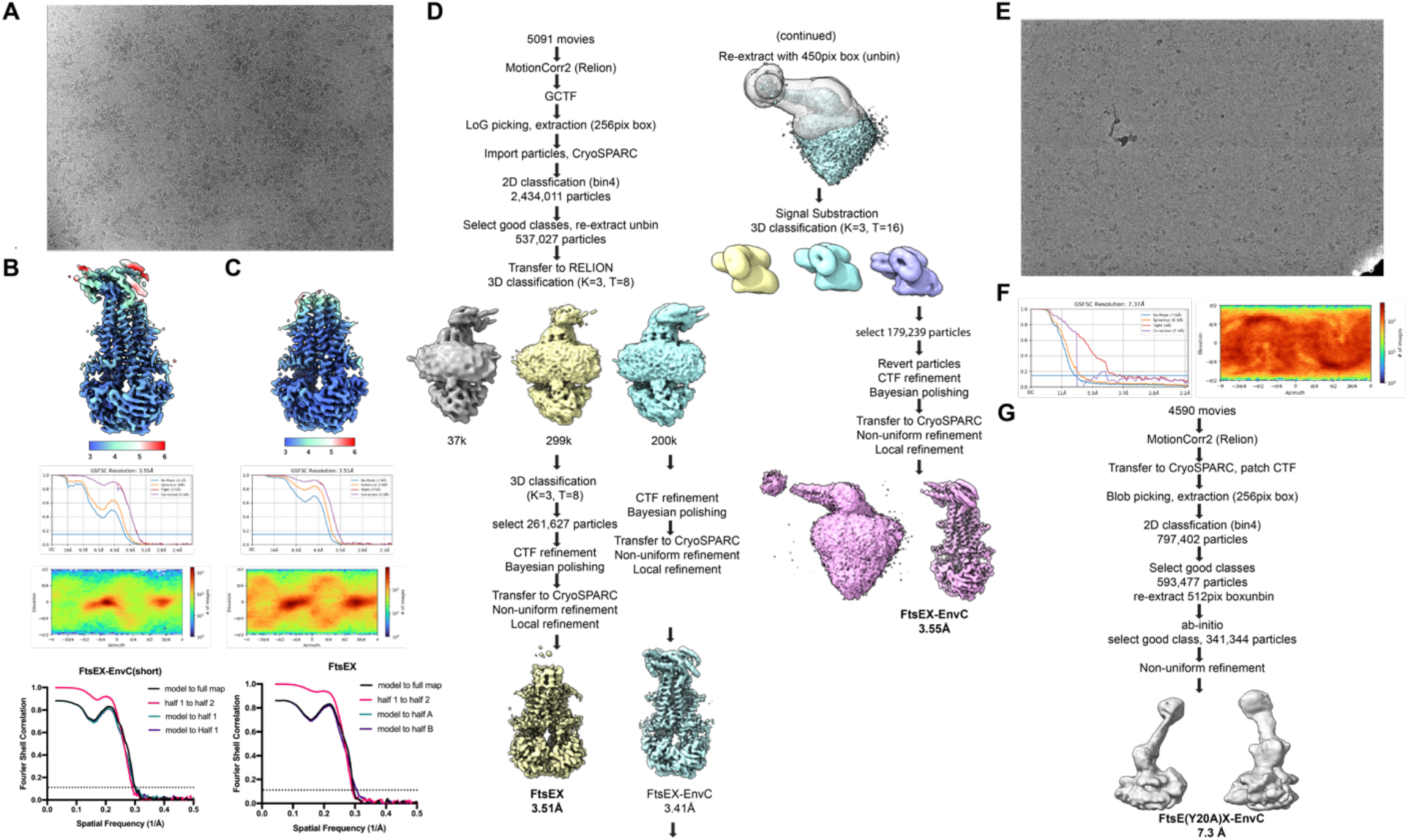
Data processing flowchart. (**A**) Representative micrograph of FtsEX-EnvC. **(B)** Local resolution estimation of FtsEX-EnvC, phenix and cryoSPARC reported Fourier shell correlations, and particle angular distribution for the final map of FtsEX-EnvC. **(C)** Local resolution estimation of FtsEX-EnvC, phenix and cryoSPARC reported Fourier shell correlations, and particle angular distribution for the final map of FtsEX. **(D)** Processing workflow for FtsEX and FtsEX-EnvC. **(E)** Representative micrograph of FtsE(Y20A)X-EnvC. **(F)** CryoSPARC reported Fourier shell correlations and particle angular distribution for the final map of FtsE(Y20A)X-EnvC. **(G)** Processing workflow for FtsEX and FtsE(Y20A)X-EnvC.

**Supplementary Figure 3.**
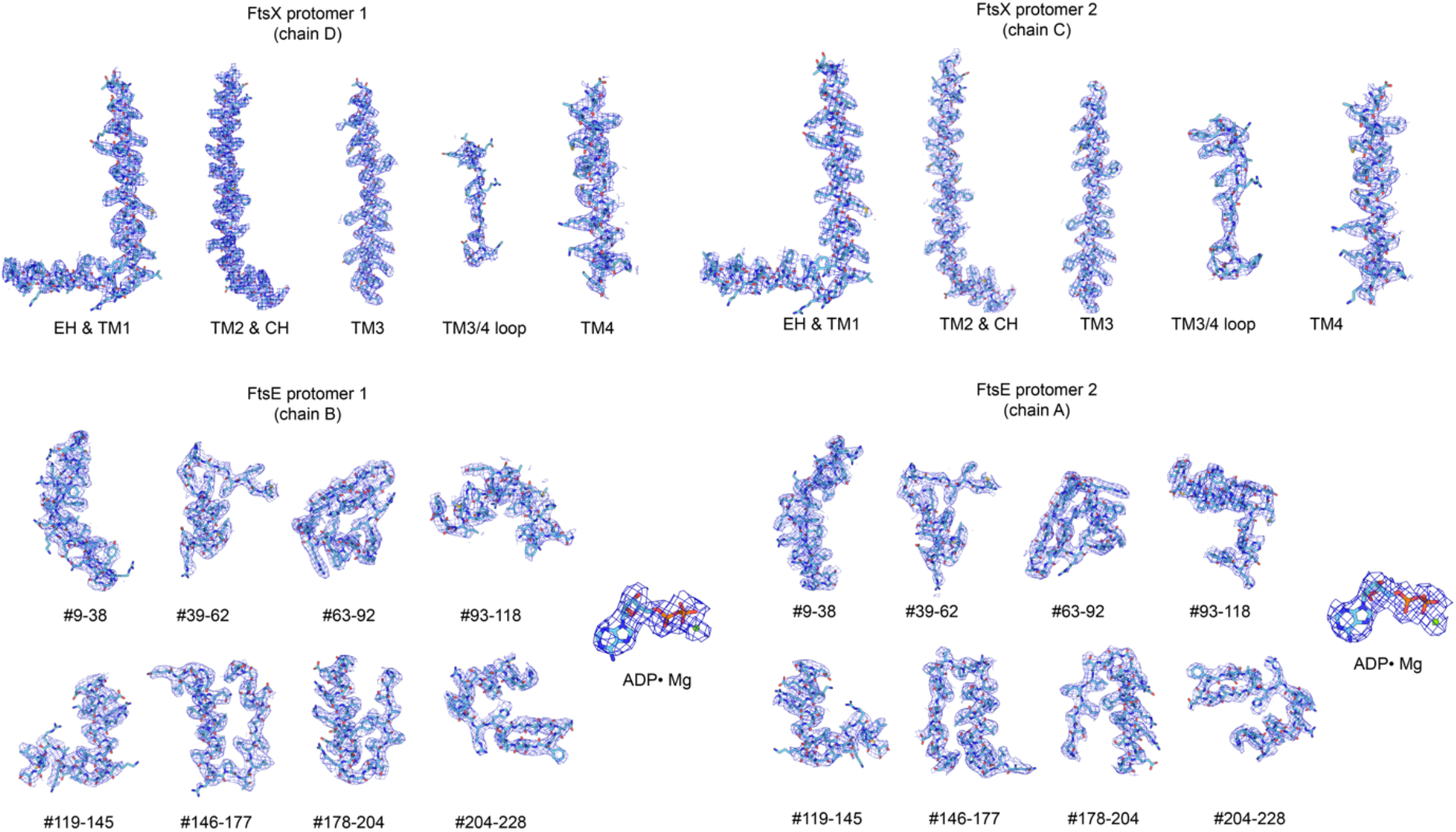
Cryo-EM density for FtsEX. Cryo-EM density corresponding to two FtsE protomers, two FtsX protomers, and ADP density.

**Supplementary Figure 4.**
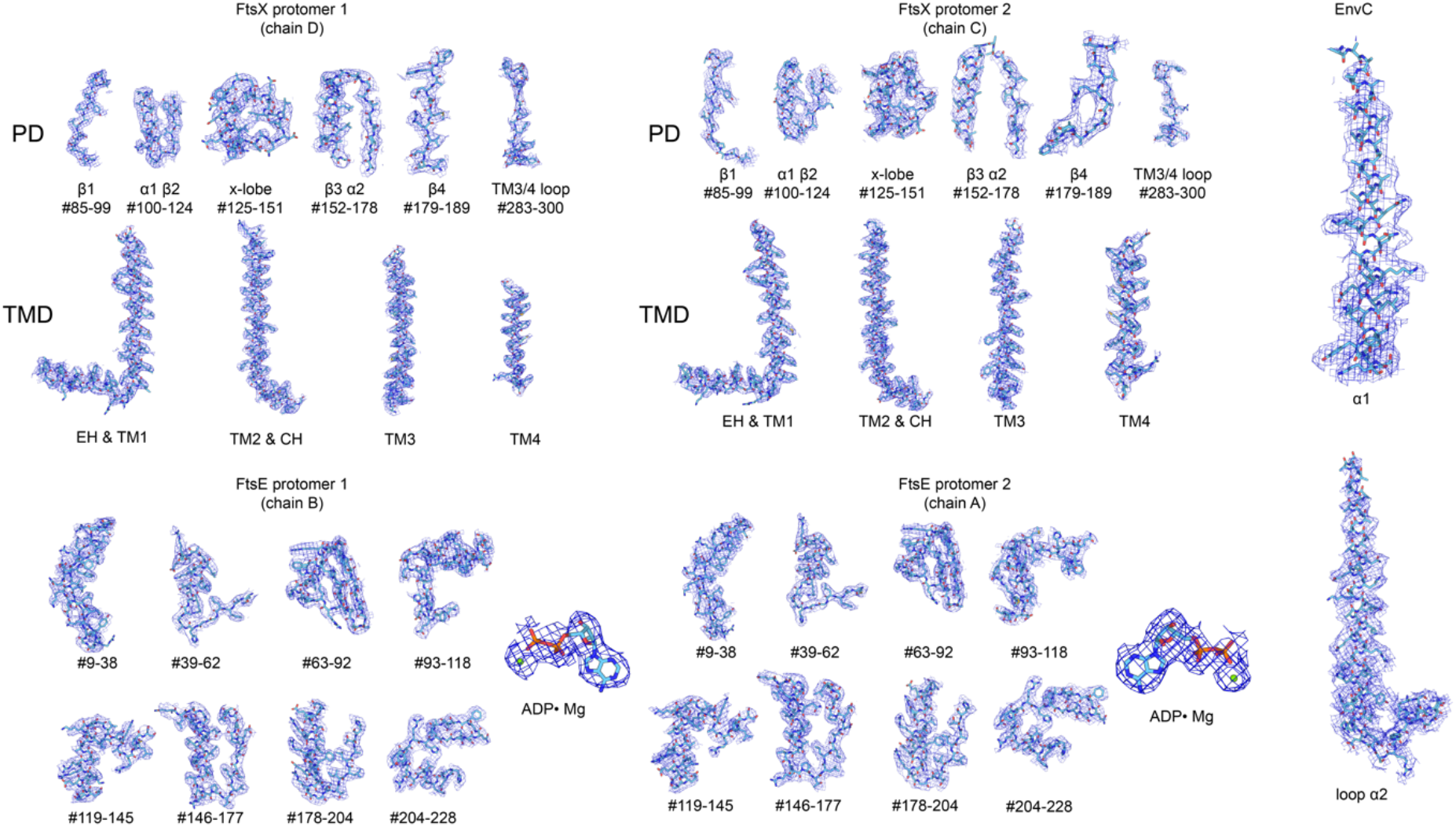
Cryo-EM density for FtsEX-EnvC. Cryo-EM density corresponding to two FtsE protomers, two FtsX protomers, EnvC and ADP density.

**Supplementary Figure 5.**
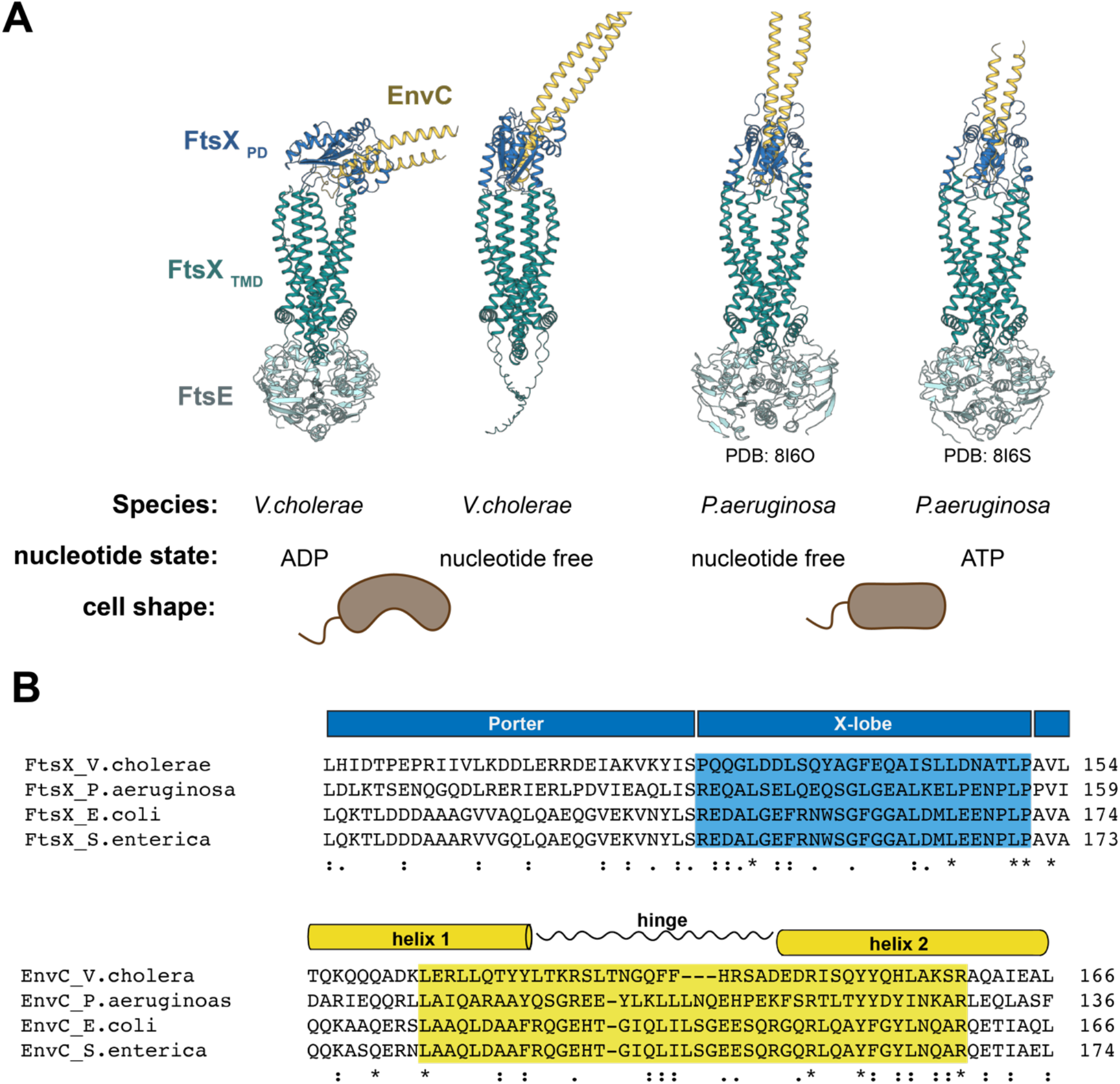
Comparison with *P.aeruginosa* FtsEX-EnvC and sequence alignment. (**A**) Structural comparison between *V.cholera* FtsEX-EnvC and *P.aeruginosa* FtsEX-EnvC. EnvC swings up and down in the *V.cholera* structures while EnvC stays vertically extended in P.aeruginosa. **(B)** Sequence alignment of FtsX and EnvC with region of inter-protein interaction sites in highlight.

